# Integrative Co-Registration of Elemental Imaging and Histopathology for Enhanced Spatial Multimodal Analysis of Tissue Sections through TRACE

**DOI:** 10.1101/2024.03.06.583819

**Authors:** Yunrui Lu, Ramsey Steiner, Serin Han, Aruesha Srivastava, Neha Shaik, Matthew Chan, Alos Diallo, Tracy Punshon, Brian Jackson, Fred Kolling, Linda Vahdat, Louis Vaickus, Jonathan Marotti, Sunita Ho, Joshua Levy

## Abstract

**Summary:** Elemental imaging provides detailed profiling of metal bioaccumulation, offering more precision than bulk analysis by targeting specific tissue areas. However, accurately identifying comparable tissue regions from elemental maps is challenging, requiring the integration of hematoxylin and eosin (H&E) slides for effective comparison. Facilitating the streamlined co-registration of Whole Slide Images (WSI) and elemental maps, TRACE enhances the analysis of tissue regions and elemental abundance in various pathological conditions. Through an interactive containerized web application, TRACE features real-time annotation editing, advanced statistical tools, and data export, supporting comprehensive spatial analysis. Notably, it allows for comparison of elemental abundances across annotated tissue structures and enables integration with other spatial data types through WSI co-registration.

**Availability and Implementation:** Available on the following platforms– GitHub: *jlevy44/trace_app*, PyPI: *trace_app*, Docker: *joshualevy44/trace_app*, Singularity: *joshualevy44/trace_app*.

**Supplementary information:** Supplementary data are available.

## 1 Implementation

There are at least twenty known trace elements that are essential for human survival, relating to energy metabolism, enzymatic activity, maintaining osmotic pressure, transport proteins, amongst others (Jomova *et al*., 2022). However, deviations from normal metal homeostasis, whether due to a deficiency or excess of these essential elements, or the introduction of toxic elements from environmental exposure, diet, or lifestyle, are linked to the development/progression of various health conditions (Mehri, 2020). For instance, in the context of cancer, elements like copper, cadmium, and iron play significant roles in mitochondrial metabolism, cell proliferation, tumorigenesis, and proangiogenic pathways (Liao *et al*., 2020). As another example, villi in the placenta mediate the transfer of nutrients and contaminants between mother and fetus (Zhang *et al*., 2015), with bioaccumulation reflecting disruptions in homeostasis (Punshon *et al*., 2019).

Metal transporter genes are essential in maintaining homeostasis through the redistribution of elements, and their conservation across species underscores their significance as a key molecular mechanism. Understanding the disruptions in metal homeostasis could unveil novel biomarkers and therapeutic targets. For example, the competitive binding of metals to these transporters can affect the bioaccumulation of toxic metals, offering a protective effect (Brooks *et al*., 2016). Additionally, copper depletion therapies have shown promise in hindering tumor migration, invasion, and metastasis (Liu *et al*., 2021; Chan *et al*., 2017; Ramchandani *et al*., 2021; Akerfeldt *et al*., 2017; Baldari *et al*., 2019; Fatfat *et al*., 2014; Gandin *et al*., 2012; Lopez *et al*., 2019; Denoyer *et al*., 2015; Gupte and Mumper, 2009). This is hypothesized to occur through two primary mechanisms: firstly, by reducing tumor metabolism (Ge *et al*., 2022; Finney *et al*., 2009; Kozono *et al*., 2018; Chellan and Sadler, 2015; Fouani *et al*., 2017), and secondly, by limiting the remodeling of the collagen extracellular matrix, which is crucial for tumor movement (Naji *et al*., 2019; Shi *et al*., 2021; Liang *et al*., 2021; Wu *et al*., 2022). Furthermore, lifestyle modifications, including dietary changes and minimizing exposure, are crucial for prevention and management.

Finding where metals are situated in tissues is a new and challenging area, similar to the task of locating genes within tissues, which is vital for unraveling the complexities of various biological systems. Traditional measurements of elemental abundance, on a bulk scale, tend to neglect the intricacies and disruptions of metal homeostasis within specific tissue architectures, obscuring critical associations and insights (Moses and Pachter, 2022). Spatially resolved metal analysis through techniques like laser ablation inductively coupled plasma time-of-flight mass spectrometry (LA-ICPTOF-MS) offers detailed maps of multi-elemental distributions at one-micron resolution (Bussweiler *et al*., 2017; Theiner *et al*., 2019; Löhr *et al*., 2019; da Silva and Arruda, 2022; Clases and Gonzalez de Vega, 2022a, 2022b; Chang *et al*., 2022). Metal ions are essential for the functioning of many biomolecules, including proteins. For instance, Zinc is an essential nutrient and absence of this element is tied to upregulation hypoxia inducible factor HIF-1α in response to oxidative stress (Choi *et al*., 2018; Marreiro *et al*., 2017). Resulting biomolecular changes may be identified with immunohistochemistry (IHC)– associated proteomic changes may localize to specific tissue architectures (Niedzwiecki *et al*., 2016). These analyses typically involve pathologists meticulously marking areas on hematoxylin and eosin (H&E) or immunohistochemical slides. Alternatively, this process can be automated through the use of computer vision technologies, such as deep learning, to annotate tissue regions and cellular phenotypes (LeCun *et al*., 2015). Deep learning algorithms excel at autonomously delineating both regions of interest within tissues and the constituent cell types, requiring minimal human oversight (Reddy *et al*., 2022). Furthermore, H&E slides are considered the gold standard for aligning with multiplex immunofluorescence (mIF), multiplexed IHC (mIHC), and other spatial molecular modalities. Such alignment offers numerous advantages for an integrative multimodal analysis, leading to a more comprehensive understanding of spatial biomolecular heterogeneity and insights into multiple mechanistic pathways.

In this paper, we introduce TRACE–Tissue Region Analysis through Co-registration of Elemental Maps. Developed by the Biomedical National Elemental Imaging Resource (BNEIR), TRACE is an interactive whole slide images (WSI) with elemental maps, encompassing various imaging formats and a range of elemental imaging techniques (e.g., LA-ICPMS, XRF). TRACE enables comparisons of metal abundance across different tissue structures and data exported from this application also can be readily integrated with additional spatial genomics assays, such as spatial transcriptomics (Ståhl *et al*., 2016; Stark *et al*., 2019; Berglund *et al*., 2018; Ji *et al*., 2020; Maynard *et al*., 2021; Fawkner-Corbett *et al*., 2021; PALISI Pediatric Intensive Care Influenza (PICFLU) Investigators *et al*., 2020; Garcia-Alonso *et al*., 2021; Meylan *et al*., 2022). TRACE allows for a streamlined data integration, preprocessing, co-registration, and analysis through the following collection of modules (**Figure 1, Supplementary Figure 1**):

**Figure 1:**
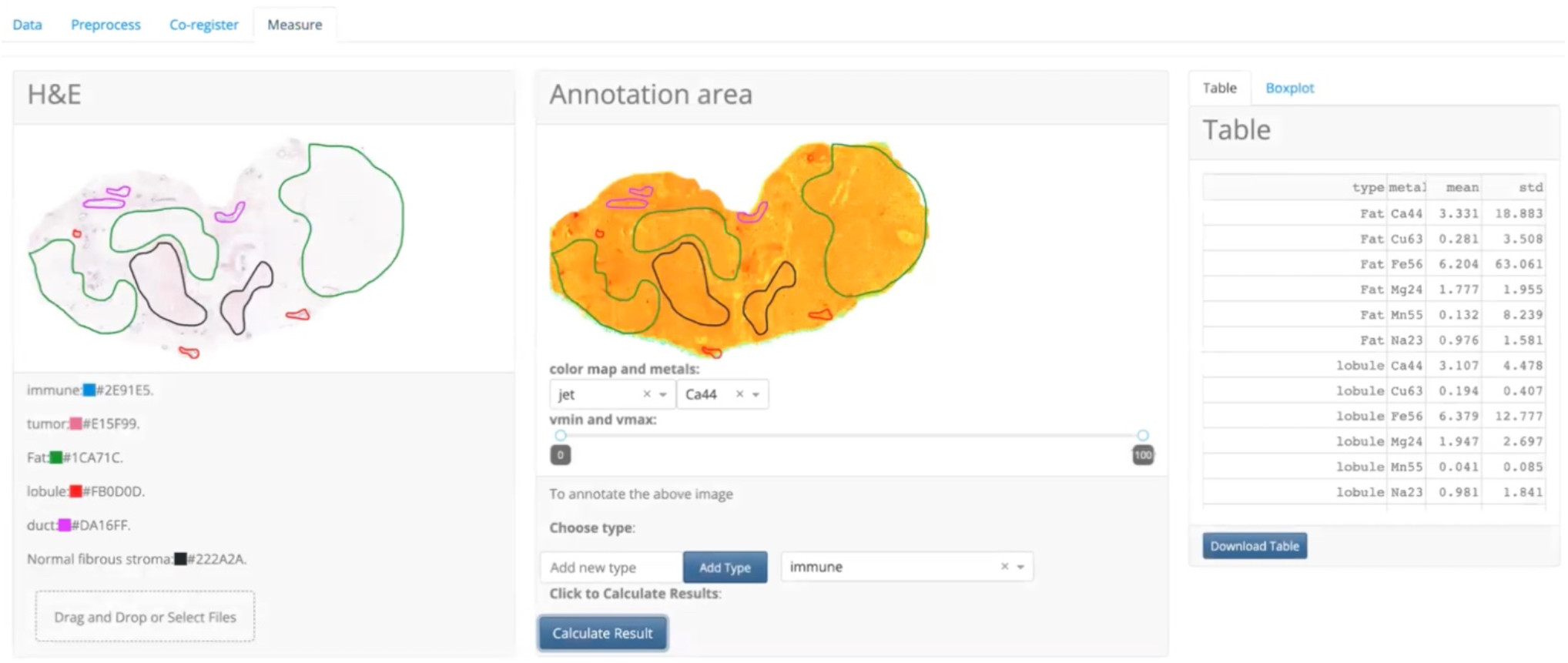
Transfer of Pathologist Annotations to LA-ICPMS Elemental Map with TRACE Co-Registration.

### 1. Efficient Data Management through Data Module

This module simplifies the organization, upload, and management of various pathology images (including H&E, IHC, mIF) and annotations in JSON/XML format from tools like QuPath and ASAP (Humphries *et al*., 2021; Bankhead *et al*., 2017; Litjens, 2017). It also handles metal images from techniques such as MALDI, LA-ICPTOF-MS, and XRF in multiple formats (Paton *et al*., 2011). The Data Module streamlines the integration of elemental imaging data, accommodating files exported from software like iolite (analyses software to generate quantitative spatial maps of elements in histologic sections, biopsies included). These files, typically a series of single channel elemental maps in Excel format, are consolidated into single files following the Bio-Format standards (e.g., OME-ZARR) for streamlined access and use (Moore *et al*., 2021).

### 2. Preprocessing and Co-Registration Modules

Once multi-channel elemental images are generated, tissue is detected using a custom workflow which aggregates elemental abundance across selected channels. This process includes smoothing of pseudo-log-transformed data using a Gaussian filter, followed by user-defined or custom thresholding and morphological operations to refine the image, such as removing extraneous objects and filling in gaps within contiguous areas. After preprocessing, co-registration takes place. While there are advanced co-registration workflows utilizing feature matching and nonlinear transformations between H&E and elemental maps, our approach is currently landmark-based. This allows for real-time labeling of similar histopathological regions of interest, enabling linear co-registration to merge H&E images with metal maps effectively. Customizable visualizations aid in enhancing landmark identification, which is vital for pathologists to correlate pathology findings with metal composition and to transfer annotated biomarkers or regions on WSI directly to elemental maps.

### 3. Data Visualization/Analysis via the Measure Module

This module facilitates the visualization of co-registered elemental maps alongside Whole Slide Images (WSI). It provides advanced tools for precise annotation and measurement within specific regions. Users can upload, import, and synchronize annotations from WSI in various formats, including JSON, XML, or GeoJSON, which are compatible with QuPath/ASAP pathology annotation tools. TRACE enables real-time synchronization between H&E and metal maps images, enhancing data interpretation, and currently provides basic statistical analysis functionalities. Additionally, data labeled with this tool can be exported in standard bioimaging formats, such as OME-ZARR, facilitating further analysis of elemental data along with the associated annotations transferred from WSI.

TRACE was implemented using a Python Dash/Flask front end (Dabbas, 2021), Python v3.9.

## 2 Results

TRACE was initially tested in two illustrative use cases, with plans for more comprehensive analyses for expanded cohorts in future works:

### 1. Annotation Transfer in Breast Cancer Study

Applied to breast tumors and normal adjacent tissue with varying HER2/HR/TNBC molecular subtypes and histologies, TRACE enabled the transfer of pathology annotations from WSI to examine architectural differences in elemental maps. The following regions were annotated by a pathologist using the QuPath annotation software: Duct, Fat, Immune Cells, Interface, Lobule, Normal Fibrous Stroma, Stroma, and Tumor. Architectures were compared if at least tumor and normal adjacent tissue were profiled as means of comparison. Preliminary results were derived using Bayesian hierarchical hurdle gamma regression models (Bürkner, 2018, 2017; Carpenter *et al*., 2017), reporting differences in tissue architectures by tissue type (tumor/normal adjacent) and architecture (**Supplementary Figure 2, Supplementary Table 1**), accounting for patient using random effects. It should be noted due to limited sample size that the returned results provide an example of what can be done and is only proof of concept and do not draw any conclusions reserved for more expansive study.

### 2. Colorectal Cancer Case Study with Spatial Transcriptomics

As a proof-of-concept, TRACE was applied to a pT3 stage colorectal cancer case, integrating with spatial transcriptomics data (M. Fatemi *et al*., 2023; M. Y. Fatemi *et al*., 2023). The 10x Genomics Visium CytAssist spatial transcriptomics (ST) assay (Janesick *et al*., 2023), capturing spatial gene expression variations within 55-micron spots, was aligned with 40X H&E-stained whole slide images (Leica Aperio GT450). Regions in and around the tumor were annotated by pathologists. ST data was clustered using UMAP embeddings and HDBSCAN and visualized, with further analysis left for a future work (**Supplementary Figure 3**) (McInnes *et al*., 2018, 2017).

Elemental imaging at 5-micron resolution utilized laser ablation inductively coupled plasma time-of-flight mass spectrometry (LA-ICPTOF-MS), on tissue sectioned from formalin-fixed, paraffin-embedded blocks. Prior to further analysis, co-registered data was exported from TRACE.

## 3 Benefits and Future Direction

TRACE permits flexible integration of histological and spatial transcriptomic data with elemental imaging analysis. This tool is available via PyPI (*trace-app*), GitHub (*jlevy44/trace_app*), Docker (*joshualevy44/trace_app*) and Singularity (*joshualevy44/trace_app*), making the tool reproducible, accessible/sharable and operating system agnostic. Applying this web application across various tissue types will broaden the scope and validity of our research in identifying prognostic elemental and transcriptomic markers within specific tissue structures. The current version allows for the co-registration of Whole Slide Images (WSI) with elemental maps. Future updates include a second release, which will integrate multiple modalities into a single multimodal data array. This approach will enable the integration of multiplexed spatial imaging and genomics assays with elemental imaging and histology. Such integration facilitates metals-based pathway analysis at a resolution approaching that of single cells. However, it is important to recognize that elemental imaging is generally a destructive process, necessitating the analysis to be conducted on consecutive tissue sections. Linking individual cell profiles with elemental abundance poses significant challenges, often requiring studies at a broader scale, such as examining the relationship between cellular interaction densities and the presence of metals and their mixtures. Furthermore, dewaxing of formalin fixed paraffin embedded tissue sections is currently recommended for LA-ICPTOF-MS applications to reduce the likelihood of signal intensity fluctuations though the impact of paraffin removal on metal distribution is currently understudied. While further spatial genomics analysis can currently be achieved by exporting and analyzing co-registered data, future enhancements will offer interactive clustering tools, identification of metal hotspots using local spatial autocorrelation statistics, advanced machine learning analytics (Paul *et al*., 2021), and enhanced nonlinear co-registration techniques. Deployment on cloud platforms like Amazon Web Services is planned to improve accessibility. Additionally, high-resolution viewing of highly multiplexed imaging will be enabled through bioimaging software like ViV, Vitessce, and Avivator, using standard NGFF formats (Manz *et al*., 2022; Keller *et al*., 2021). The integration of these features, which currently require external analysis of exported data, will further enhance the application’s capabilities.

## Funding

This work has been supported by the NIH grants R24GM141194, P20GM104416, and P20GM130454 subawards to JL.

### Conflict of Interest

None declared.

## Notes

### Competing Interest Statement

The authors have declared no competing interest.

