## Supplementary Materials for "Integrative Co-Registration of Elemental Imaging and Histopathology for Enhanced Spatial Multimodal Analysis of Tissue Sections through TRACE"

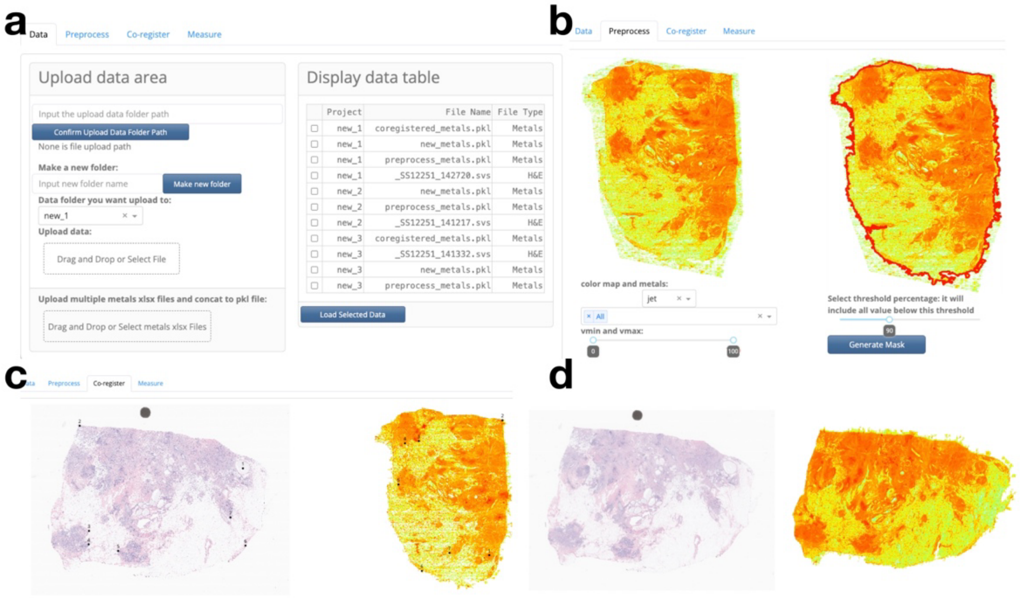


**Supplementary Figure 1: Demonstration of TRACE Web Application Components: a)** Data upload and management module selects H&E whole slide image and elemental map, **b)** Preprocessing module identifies tissue mask to remove background signal, **c)** Landmarks identified between H&E WSI and LA-ICPMS image to facilitate co-registration, **d)** Annotations are transferred from H&E section to co-registered elemental map for further measurement and downstream analysis

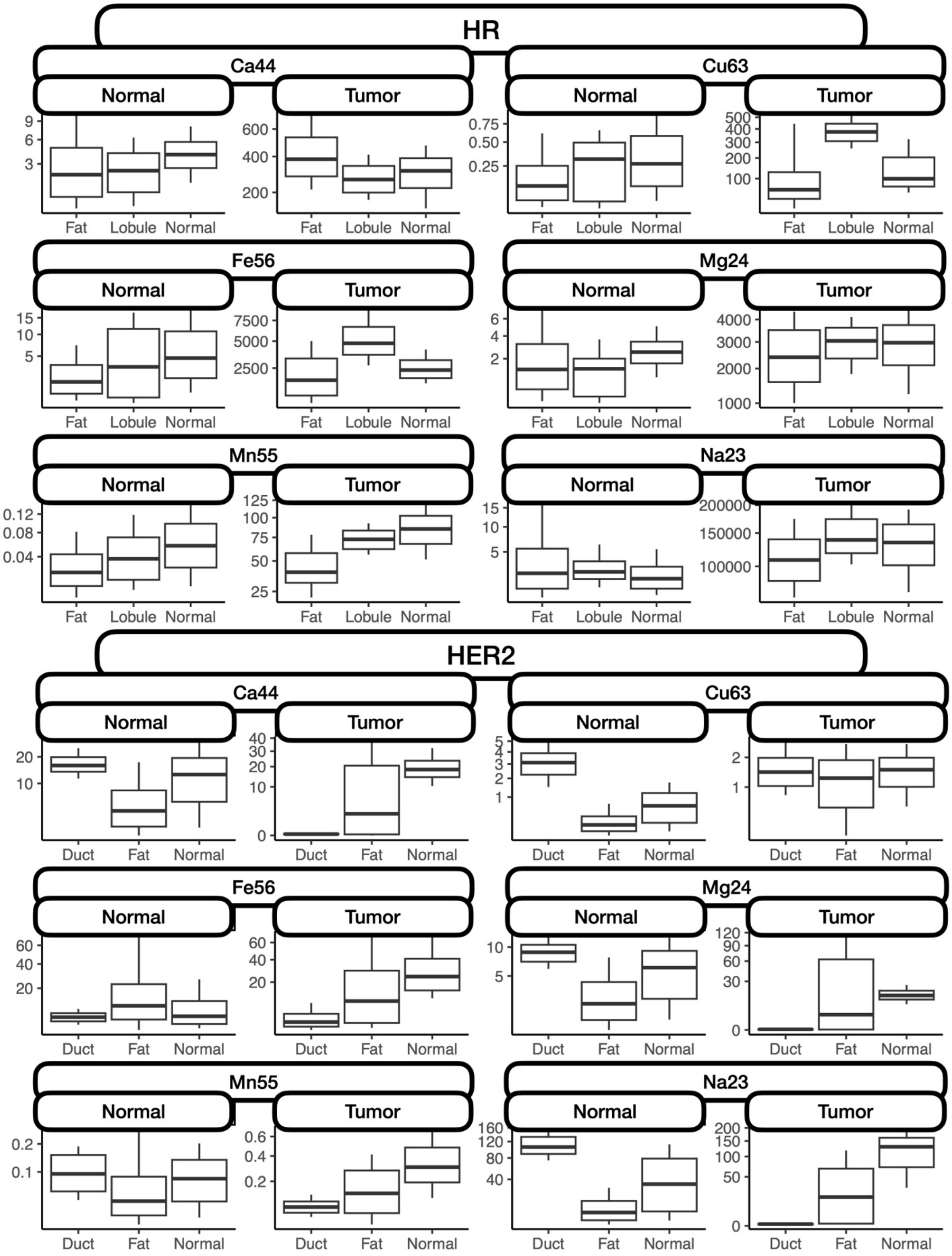
HR
Ca44
Normal
Tumor
9
6
3
600
400
200
Fat
Lobule
Normal
Cu63
0.75
0.50
0.25
500
400
300
200
100
Fe56
15
10
5
7500
5000
2500
Mg24
6
4
2
4000
3000
2000
1000
Mn55
0.12
0.08
0.04
125
100
75
50
25
Na23
15
10
5
200000
150000
100000
HER2
Ca44
20
10
40
30
20
10
0
Duct
Fat
Normal
Cu63
5
4
3
2
1
2
1
Fe56
60
40
20
60
40
20
Mg24
10
5
120
90
60
30
0
Mn55
0.2
0.1
0.6
0.4
0.2
Na23
160
120
80
40
200
150
100
50
0

**Supplementary Figure 2: Boxplot comparison of elemental abundance in various tissue regions within select breast tumor sections, comparing ducts, fat and normal adjacent tissue**

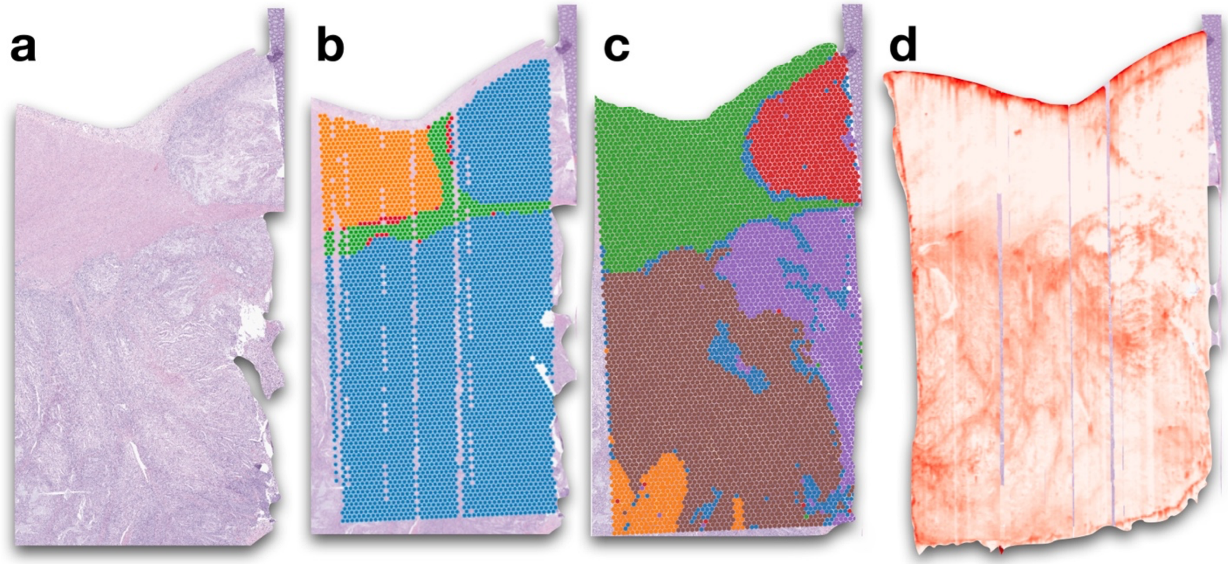


**Supplementary Figure 3: Co-Registration with TRACE on select Colorectal tumor section facilitates metals-based pathway assessment integrating: a)** H&E WSI, with **b)** pathologist annotations to delineate tissue architectures, **C)** spatial transcriptomics data, expression of nearly 18,000 genes at 55-micron resolution and **D)** elemental imaging maps with LA-ICPTOF-MS at 5-micron resolution for all elements of the periodic table

### Supplementary Table 1: Statistical Analysis Results for LA-ICPMS-based differential elemental abundance between various tissue architectures for select specimens reflecting breast tumor subtypes

| Subtype | Element | Parameter | Coefficient | 2.5% | 97.5% | p-value |
|---|---|---|---|---|---|---|
| **HER2** | **Ca44** | Duct > Fat, N | 0.22 | 0.19 | 0.26 | <0.001 |
| | | Duct > Normal Fibrous Stroma, N | 0.08 | 0.04 | 0.12 | <0.001 |
| | | Fat > Normal Fibrous Stroma, N | -0.14 | -0.20 | -0.09 | <0.001 |
| | | Duct > Fat, T | -0.42 | -0.47 | -0.38 | <0.001 |
| | | Duct > Normal Fibrous Stroma, T | -0.06 | -0.14 | 0.02 | 0.18 |
| | | Fat > Normal Fibrous Stroma, T | 0.37 | 0.27 | 0.46 | <0.001 |
| | **Cu63** | Duct > Fat, N | 2.16 | 2.13 | 2.19 | <0.001 |
| | | Duct > Normal Fibrous Stroma, N | 1.55 | 1.52 | 1.59 | <0.001 |
| | | Fat > Normal Fibrous Stroma, N | -0.61 | -0.65 | -0.56 | <0.001 |
| | | Duct > Fat, T | 0.08 | 0.04 | 0.12 | <0.001 |
| | | Duct > Normal Fibrous Stroma, T | -0.53 | -0.60 | -0.46 | <0.001 |
| | | Fat > Normal Fibrous Stroma, T | -0.61 | -0.70 | -0.53 | <0.001 |
| | **Fe56** | Duct > Fat, N | -1.90 | -1.95 | -1.84 | <0.001 |
| | | Duct > Normal Fibrous Stroma, N | -0.85 | -0.91 | -0.79 | <0.001 |
| | | Fat > Normal Fibrous Stroma, N | 1.05 | 0.97 | 1.13 | <0.001 |
| | | Duct > Fat, T | -1.39 | -1.46 | -1.32 | <0.001 |
| | | Duct > Normal Fibrous Stroma, T | -2.65 | -2.77 | -2.54 | <0.001 |
| | | Fat > Normal Fibrous Stroma, T | -1.26 | -1.41 | -1.12 | <0.001 |
| | **Mg24** | Duct > Fat, N | -0.56 | -0.61 | -0.51 | <0.001 |
| | | Duct > Normal Fibrous Stroma, N | -1.11 | -1.16 | -1.06 | <0.001 |
| | | Fat > Normal Fibrous Stroma, N | -0.55 | -0.62 | -0.48 | <0.001 |
| | | Duct > Fat, T | -2.75 | -2.81 | -2.68 | <0.001 |
| | | Duct > Normal Fibrous Stroma, T | -1.60 | -1.71 | -1.50 | <0.001 |
| | | Fat > Normal Fibrous Stroma, T | 1.14 | 1.01 | 1.27 | <0.001 |
| | **Mn55** | Duct > Fat, N | -1.42 | -1.47 | -1.37 | <0.001 |
| | | Duct > Normal Fibrous Stroma, N | -2.41 | -2.45 | -2.35 | <0.001 |
| | | Fat > Normal Fibrous Stroma, N | -0.99 | -1.06 | -0.92 | <0.001 |
| | | Duct > Fat, T | -0.42 | -0.48 | -0.36 | <0.001 |
| | | Duct > Normal Fibrous Stroma, T | -1.08 | -1.19 | -0.97 | <0.001 |
| | | Fat > Normal Fibrous Stroma, T | -0.66 | -0.79 | -0.53 | <0.001 |
| | **Na23** | Duct > Fat, N | 0.90 | 0.87 | 0.93 | <0.001 |
| | | Duct > Normal Fibrous Stroma, N | 0.64 | 0.61 | 0.67 | <0.001 |
| | | Fat > Normal Fibrous Stroma, N | -0.26 | -0.30 | -0.21 | <0.001 |
| | | Duct > Fat, T | -0.24 | -0.28 | -0.20 | <0.001 |
| | | Duct > Normal Fibrous Stroma, T | 0.32 | 0.26 | 0.39 | <0.001 |
| | | Fat > Normal Fibrous Stroma, T | 0.57 | 0.48 | 0.64 | <0.001 |
| **HR** | **Ca44** | Fat > Lobule, N | 0.71 | 0.58 | 0.85 | <0.001 |
| | | Fat > Normal Fibrous Stroma, N | 0.21 | 0.17 | 0.25 | <0.001 |
| | | Lobule > Normal Fibrous Stroma, N | -0.51 | -0.65 | -0.36 | <0.001 |
| | | Fat > Lobule, T | 0.37 | 0.12 | 0.59 | 0.004 |
| | | Fat > Normal Fibrous Stroma, T | 0.14 | 0.02 | 0.26 | 0.022 |
| | | Lobule > Normal Fibrous Stroma, T | -0.22 | -0.48 | 0.05 | 0.094 |
| | **Cu63** | Fat > Lobule, N | 0.15 | 0.03 | 0.25 | 0.008 |
| | | Fat > Normal Fibrous Stroma, N | -0.09 | -0.13 | -0.05 | <0.001 |
| | | Lobule > Normal Fibrous Stroma, N | -0.24 | -0.36 | -0.11 | <0.001 |
| | | Fat > Lobule, T | -0.60 | -0.80 | -0.42 | <0.001 |
| | | Fat > Normal Fibrous Stroma, T | 0.33 | 0.23 | 0.43 | <0.001 |
| | | Lobule > Normal Fibrous Stroma, T | 0.93 | 0.73 | 1.16 | <0.001 |
| | **Fe56** | Fat > Lobule, N | -0.06 | -0.18 | 0.06 | 0.358 |
| | | Fat > Normal Fibrous Stroma, N | -0.22 | -0.25 | -0.18 | <0.001 |
| | | Lobule > Normal Fibrous Stroma, N | -0.16 | -0.28 | -0.03 | 0.012 |
| | | Fat > Lobule, T | -1.47 | -1.68 | -1.26 | <0.001 |
| | | Fat > Normal Fibrous Stroma, T | -0.56 | -0.65 | -0.47 | <0.001 |
| | | Lobule > Normal Fibrous Stroma, T | 0.91 | 0.68 | 1.15 | <0.001 |
| | **Mg24** | Fat > Lobule, N | 0.36 | 0.24 | 0.47 | <0.001 |
| | | Fat > Normal Fibrous Stroma, N | -0.14 | -0.17 | -0.10 | <0.001 |
| | | Lobule > Normal Fibrous Stroma, N | -0.50 | -0.62 | -0.37 | <0.001 |
| | | Fat > Lobule, T | -1.16 | -1.37 | -0.96 | <0.001 |
| | | Fat > Normal Fibrous Stroma, T | -1.03 | -1.11 | -0.93 | <0.001 |
| | | Lobule > Normal Fibrous Stroma, T | 0.14 | -0.08 | 0.37 | 0.208 |
| | **Mn55** | Fat > Lobule, N | 0.34 | 0.20 | 0.48 | <0.001 |
| | | Fat > Normal Fibrous Stroma, N | -1.47 | -1.51 | -1.42 | <0.001 |
| | | Lobule > Normal Fibrous Stroma, N | -1.81 | -1.96 | -1.66 | <0.001 |
| | | Fat > Lobule, T | 0.98 | 0.74 | 1.22 | <0.001 |
| | | Fat > Normal Fibrous Stroma, T | 0.74 | 0.59 | 0.89 | <0.001 |
| | | Lobule > Normal Fibrous Stroma, T | -0.24 | -0.51 | 0.05 | 0.088 |
| | **Na23** | Fat > Lobule, N | -0.30 | -0.42 | -0.19 | <0.001 |
| | | Fat > Normal Fibrous Stroma, N | -0.24 | -0.28 | -0.21 | <0.001 |
| | | Lobule > Normal Fibrous Stroma, N | 0.06 | -0.06 | 0.19 | 0.344 |
| | | Fat > Lobule, T | -0.64 | -0.85 | -0.43 | <0.001 |
| | | Fat > Normal Fibrous Stroma, T | -0.41 | -0.51 | -0.31 | <0.001 |
| | | Lobule > Normal Fibrous Stroma, T | 0.23 | 0.00 | 0.47 | 0.054 |
